# Capsule types driving carbapenem-resistant *Klebsiella pneumoniae* in Pakistan are largely absent from the phage literature

**DOI:** 10.64898/2026.09.29.755075

**Authors:** Talal Ahmed, Ahmed Hussain Shah

## Abstract

Phage therapy is increasingly proposed for carbapenem-resistant *Klebsiella pneumoniae*, but most *Klebsiella* phages bind a single capsule type, so a phage is useful only where its capsule type circulates. We asked whether published phages cover the capsule types found in Pakistani patients.

We typed 254 clinical *K. pneumoniae* genomes from Pakistan and curated the phage and depolymerase literature into 34 phage–capsule records covering 27 phages and 15 capsule types. Nearly half the cohort was unmatched: 121 of 254 isolates (47.6%), spanning 51 of 66 capsule types, belong to types for which no phage has been described, and 67 of those isolates carry a carbapenemase. Collapsing each BioProject to one isolate per type raises the unmatched fraction to 51.8%.

To ask why, we typed 475 genomes sampled on identical criteria from India, Bangladesh and Nepal, China, Europe and the USA. Two capsule types separated sharply. KL81 accounted for 6.3% of South Asian genomes and none of 275 from China, Europe or the USA, and has no published phage; KL107 showed the reverse, 9.8% in those regions and absent from South Asia. Both survived collapsing (*P* = 0.0014 and *P* = 0.0002). Six further differences did not, and are attributable to clonal expansion within single studies.

Phages are isolated against locally available hosts, so the capsule types entering the global phage collection track the regions doing the isolating. KL81 is a defined target for phage isolation in South Asia.

**Impact Statement:** Phage therapy is one of the few remaining options for carbapenem-resistant *Klebsiella pneumoniae*, and because most *Klebsiella* phages recognise a single capsule type, a cocktail only helps where its capsule types circulate. This study shows that almost half of Pakistani clinical isolates belong to capsule types for which no phage has ever been described, and that the gap is not random: the uncovered types are South Asian ones that are rare or absent in the countries where *Klebsiella* phages are isolated. The global phage collection reflects the capsule distribution of the regions doing the sampling. The practical consequence is that imported cocktails are unlikely to fit South Asian patients, and that KL81 — absent from 275 genomes from China, Europe and the USA, and carbapenemase-positive in every Pakistani isolate examined — is a specific, justified target for local phage isolation.

**Data Summary:**

1. All genome data analysed are publicly available assemblies from NCBI.
2. Pakistani cohort: 254 *Klebsiella pneumoniae* assemblies, accessions listed in Table S1.
3. Comparison cohort: 475 assemblies from India, Bangladesh, Nepal, China, Europe and the USA, accessions listed in Table S2.
4. Source metadata: NCBI Pathogen Detection release PDG000000012.2494, https://www.ncbi.nlm.nih.gov/pathogens/
5. Curated phage–capsule evidence table (34 records, 27 phages, 15 capsule types) with PMIDs and DOIs: Table S3.
6. Capsule typing performed with Kleborate v3.2.4 and Kaptive v3.2.2, both openly available (https://github.com/klebgenomics).
7. Analysis code: https://github.com/wwwtalalahmed98-collab/phagematch-pk, archived at Zenodo, doi:10.5281/zenodo.22025259 (concept DOI, resolving to the most recent version; the version analysed here is v1.0.0, doi:10.5281/zenodo.22025260).

No new sequence data were generated by this study.

## Introduction

Carbapenem resistance in *Klebsiella pneumoniae* has moved from a rarity to a routine finding in South Asian hospitals, and the drugs held in reserve for it are running out. Phage therapy is one of the few options that does not depend on new chemistry. Individual patients have been treated successfully with tailored preparations [1, 2], and interest in assembling defined cocktails against carbapenem-resistant *K. pneumoniae* (CRKP) has grown accordingly.

The obstacle is specificity. *K. pneumoniae* is wrapped in a polysaccharide capsule, and for most lytic *Klebsiella* phages that capsule is the receptor. Phages carry tail-spike depolymerases that cleave one capsular polysaccharide and not others, so host range usually tracks capsule type closely [3]. The Kaptive v3.2.2 reference database used here contains 161 distinct *Klebsiella* capsule (K) loci [4, 5]. A phage raised against a KL64 strain is generally useless against a KL81 one.

That makes phage therapy a matching problem before it is a manufacturing problem. A cocktail is only as good as the overlap between the capsule types it covers and the capsule types infecting patients. Phage discovery is also geographically concentrated: most characterised *Klebsiella* phages and depolymerases come from China, Europe and the United States, and phages are isolated by screening environmental samples against whatever host strains are at hand. Whether the resulting collection matches the capsule types circulating in South Asia has not, to our knowledge, been tested.

We asked three questions. Which capsule types occur in Pakistani clinical *K. pneumoniae*? Which have a published phage or depolymerase? And where the answer is none, is that because the type is globally rare, or because it is common in South Asia and rare where phages are made?

## Methods

### Genome selection

Pakistani *K. pneumoniae* genomes were drawn from NCBI Pathogen Detection (release PDG000000012.2494). Isolates were retained if labelled *Klebsiella pneumoniae*, with `epi_type = clinical`, a collection date of 2015 or later, and an assembly accession. This yielded 260 genomes; six returned a Kaptive `Untypeable` capsule call and were excluded, leaving 254 for analysis.

The comparison cohort was assembled from the same metadata release under identical criteria, sampling 100 genomes each from India, Bangladesh with Nepal, China, nine western European countries, and the USA, using a fixed random seed (20260815). Of 500 accessions, 475 remained retrievable and were typed; 466 gave a typeable capsule call.

### Capsule typing

Genomes were typed with Kleborate v3.2.4 [6] using the `kpsc` preset, which calls capsule (K) and O loci through Kaptive [4, 5]. Kaptive was pinned to v3.2.2; v3.3.0 restructured the package and removed the `kaptive.database` module that Kleborate’s capsule module imports. Calls returned by Kaptive as `Untypeable` were excluded throughout — six of 260 Pakistani and nine of 475 comparison genomes.

Typing ran in chunks of 20 genomes with per-chunk completion markers written to persistent storage, allowing an interrupted run to resume from the last finished chunk. Genomes were processed in shuffled rather than sorted accession order: accession numbers cluster by submitting centre, so an interrupted sorted run leaves a geographically skewed subset, whereas any prefix of a shuffled run remains a balanced random sample across groups.

### Literature curation

PubMed was searched for `Klebsiella[tiab] AND depolymerase[tiab]` (129 records), supplemented by targeted searches for every capsule type with three or more isolates in the Pakistani cohort. Records were read and reduced to a table of phage–capsule pairings graded by evidence:

- **depolymerase characterised** — enzyme expressed and shown to degrade that capsule
- **structure solved** — as above, with a solved structure
- **phage host confirmed** — phage lyses strains of the type, no enzyme characterised
- **genome announcement only**

Host-range reports were counted as evidence for a capsule type only where the host was assigned to a single type. Reports typing hosts by *wzi* sequencing alone were not counted when the allele is compatible with several K types: the common allele *wzi*50, for example, is shared by K15, K17, K50, K51 and K52, so lysis of a *wzi*50 host does not establish activity against any one of them. Applying this rule excludes at least one recent phage-cocktail study whose abstract lists those five types together.

Curation produced 34 records covering 27 phages and 15 capsule types, drawn from the 27 primary reports cited as [8–34] and listed individually with evidence grade in Table S3. Eleven types have a characterised depolymerase; four (KL17, KL24, KL47, KL112) rest on host-range evidence alone.

### Statistics

Capsule frequencies were compared with two-sided Fisher exact tests, computed by summing the hypergeometric tail. Every capsule type with at least four isolates across the two comparison cohorts was tested, giving 31 tests, so a Bonferroni-corrected threshold of α = 0.05/31 = 0.0016 is applied throughout. All 31 are reported in Table S5.

Public genome collections are assembled largely from outbreak investigations, so genomes sharing a BioProject may represent one transmission chain rather than independent samples. Every comparison was run twice: on all genomes, and with each BioProject collapsed to a single genome per capsule type (South Asia n = 140, phage-source regions n = 212). Only results surviving the collapse are reported as findings.

## Results

### Capsule diversity in Pakistani clinical *K. pneumoniae*

The 254 Pakistani isolates carry 66 distinct capsule types, a mean of 3.8 isolates per type, and 27 types appear in a single patient. The commonest type, KL64, accounts for 9.8% of isolates and the six commonest together for 36% (Fig. 1). No type dominates, which sets a floor on how few phages a useful cocktail could contain.

**Fig. 1.**
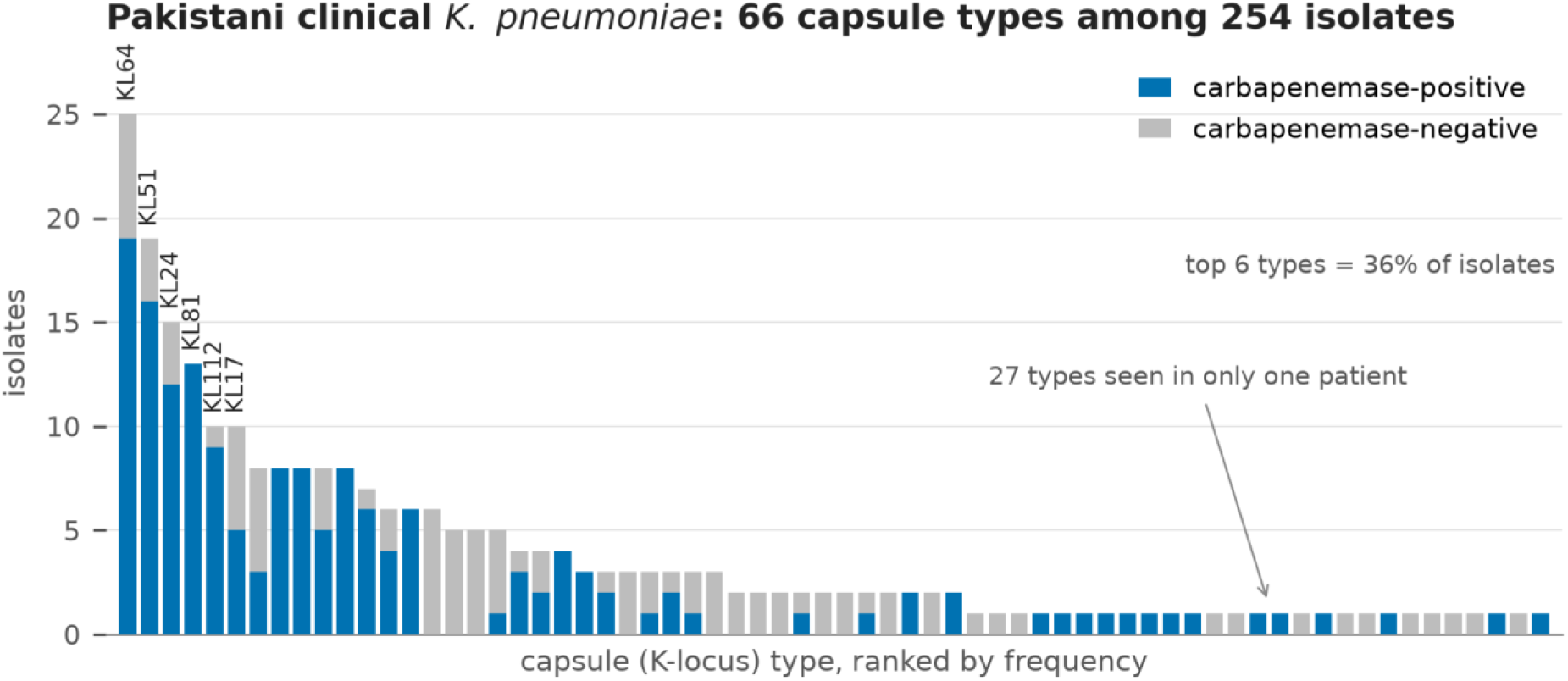
Rank-abundance of all 66 capsule (K) loci among 254 Pakistani clinical *K. pneumoniae* isolates, each bar divided into carbapenemase-positive and carbapenemase-negative isolates. The complete distribution is shown rather than a top-N subset; 27 types occur in a single patient.

### Published phage coverage

Joining the curated literature to the cohort leaves 121 of 254 isolates (47.6%), across 51 of 66 capsule types, with no published phage of any kind. Sixty-seven of those isolates carry a carbapenemase. The uncovered fraction is not a tail of singletons: KL81 (13 isolates), KL10 (8), KL36 (8) and KL15 (8) account for 37 of them, and the first three are carbapenemase-positive in every isolate (Fig. 2). Collapsing BioProjects moves the uncovered fraction to 51.8% (71 of 137), so the headline figure is conservative.

**Fig. 2.**
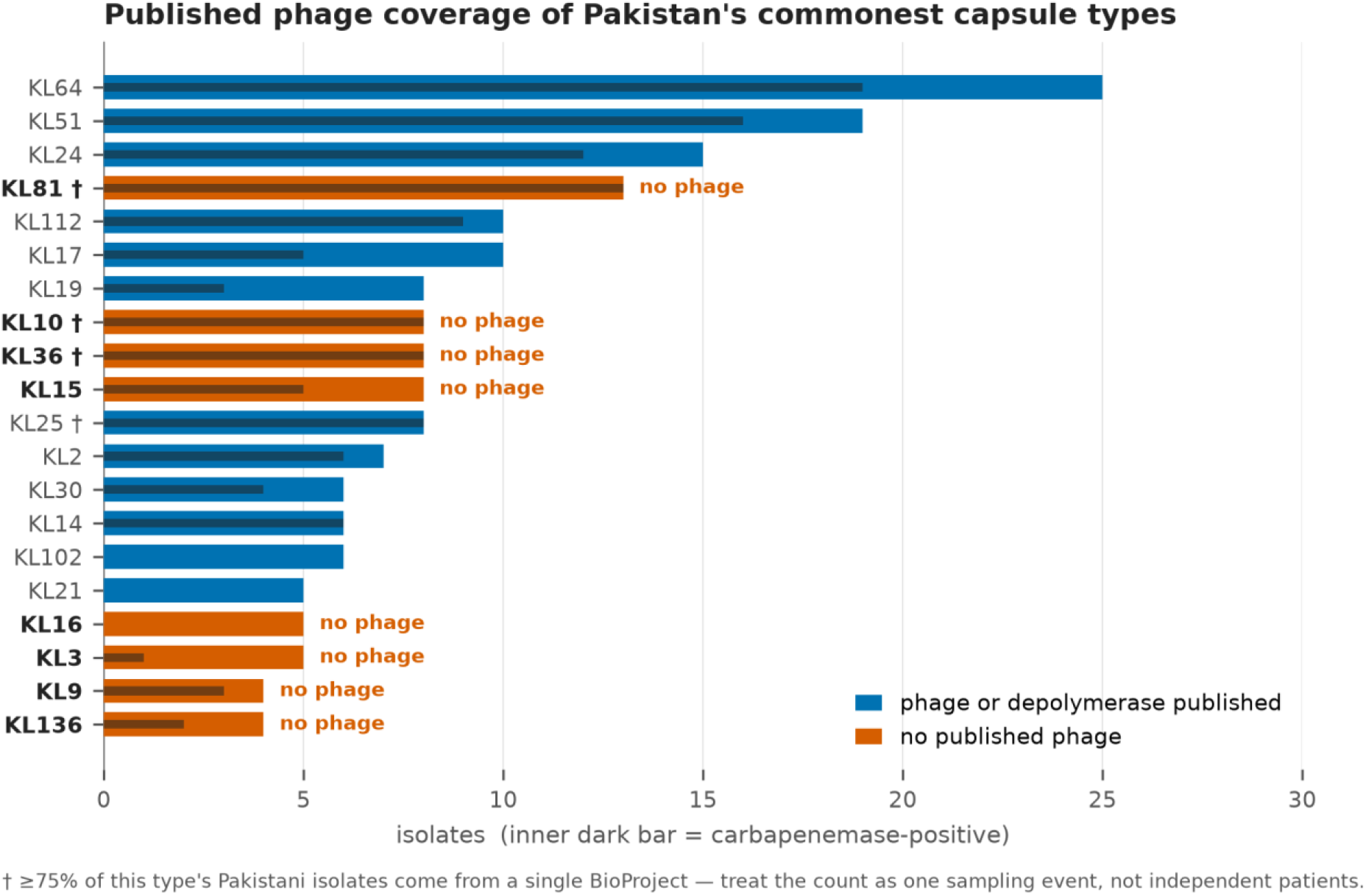
The 20 commonest capsule types, coloured by whether any phage or depolymerase has been published, with the inner dark bar showing carbapenemase-positive isolates on the same scale. Daggered types draw ≥75% of their Pakistani isolates from a single BioProject.

These per-type counts describe sampling events rather than independent patients. All 13 Pakistani KL81 genomes come from a single BioProject, as do 6 of 8 KL10 and 6 of 8 KL36 genomes, and two projects supply 61% of the cohort. The data therefore support no claim about how common any capsule type is within Pakistan.

### Capsule types differ between South Asia and the phage-producing regions

Two capsule types separate sharply between South Asia and the regions producing most characterised *Klebsiella* phages, and both survive collapsing (Fig. 3, Table 1).

**Fig. 3.**
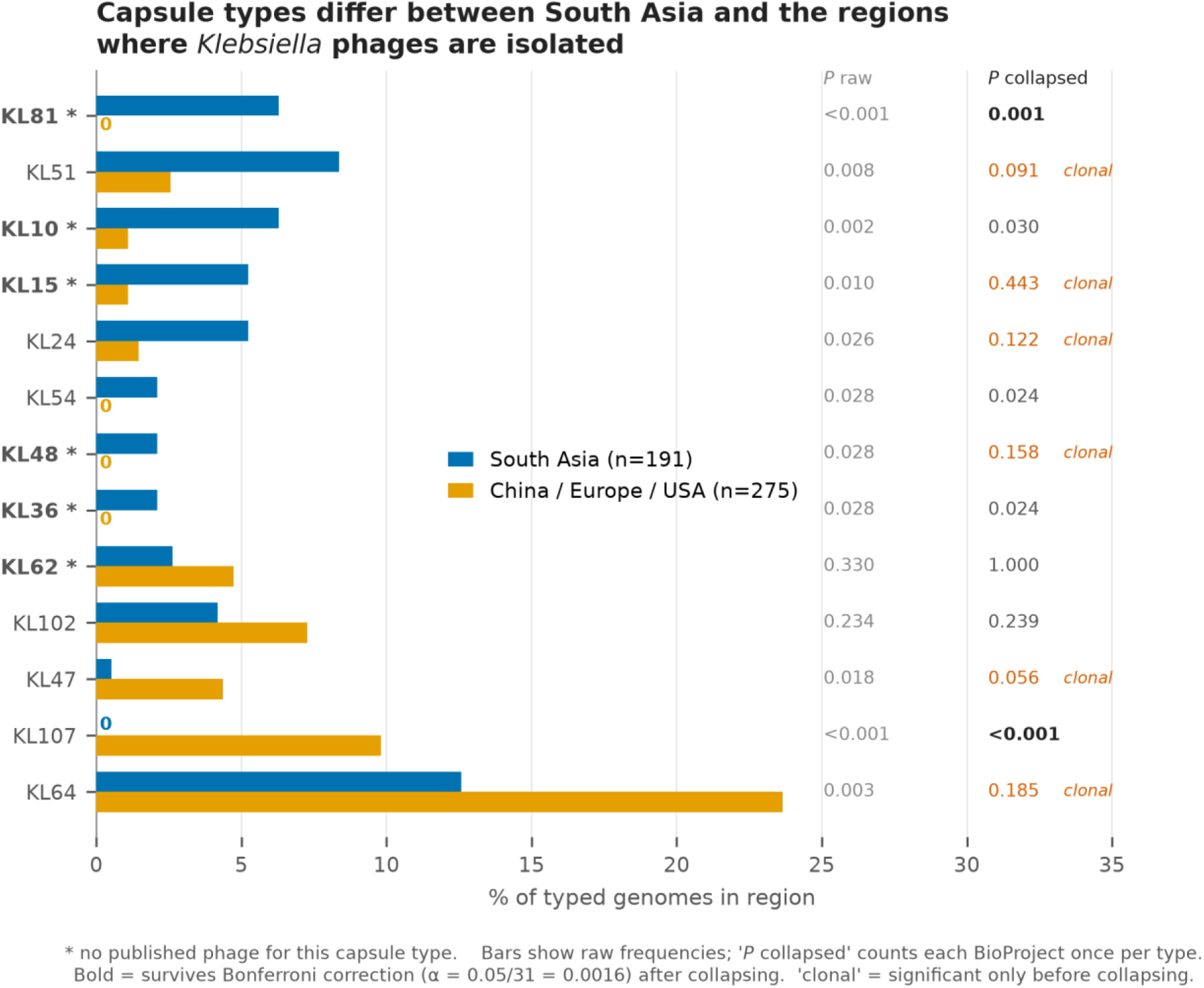
Capsule frequency in South Asia (India, Bangladesh, Nepal; n = 191) versus China, Europe and the USA (n = 275), ordered by difference. Types with no published phage are starred. Two *P*-value columns are given: raw, and with each BioProject collapsed to one genome per capsule type. Only KL81 and KL107 survive collapsing under Bonferroni correction.

**Table 1.**
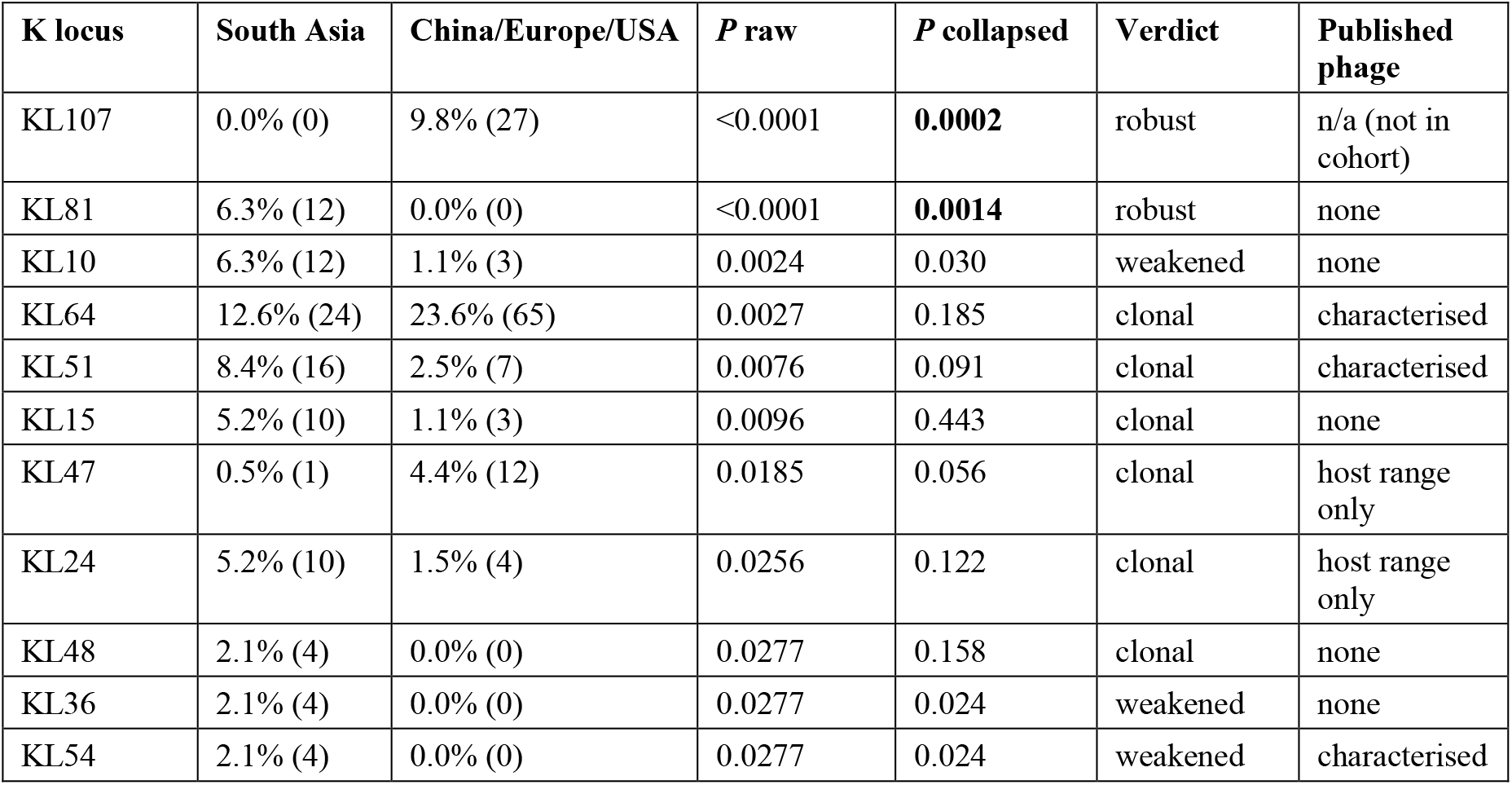
Capsule frequency by region, raw and BioProject-collapsed. All eleven capsule types reaching *P* < 0.05 on raw counts are shown, ordered by raw *P*; the remaining twenty tested types are in Table S5. Percentages are of typed genomes in each region, with isolate counts in parentheses (South Asia = India, Bangladesh and Nepal, *n* = 191; China/Europe/USA *n* = 275; the Pakistani cohort is separate and not included here).

All twelve KL81 genomes were South Asian (Bangladesh/Nepal 7, India 5) and none occurred in China, Europe or the USA. These twelve derive from seven distinct BioProjects across two countries, so the pattern is not a single outbreak: it survives collapsing (7 vs 0, *P* = 0.0014) and Bonferroni correction. KL107 is the mirror image and equally robust, drawn from 17 distinct BioProjects.

The KL81 result held as the comparison set grew. An interim analysis at 435 genomes gave 10 vs 0; the additional 40 genomes added two further South Asian KL81 isolates and none elsewhere.

Six differences — KL15, KL24, KL47, KL48, KL51 and the apparent depletion of KL64 in South Asia — were significant on raw counts and lost significance on collapsing (*P* = 0.443, 0.122, 0.056, 0.158, 0.091 and 0.185). They are attributable to clonal expansion and are not reported as findings.

## Discussion

The defensible result is narrow: KL81 is a South Asian capsule type, it does not appear in 275 genomes from China, Europe and the USA, and no phage or depolymerase against it has been published. KL107 is its mirror image. Everything else in the comparison weakened or collapsed once clonal structure was accounted for.

That narrow result carries a general point. Phages are found by screening sewage, hospital effluent or water against host strains the finding laboratory already holds. The capsule types entering the world’s phage collection are therefore a sample of the capsule types available to the laboratories doing the sampling. When those laboratories are concentrated in a few countries, the collection inherits their local capsule distribution. KL81 was never in the water being screened, and the absence of a KL81 phage follows from where the screening happened rather than from any oversight.

For practice this reframes the problem. There is no shortage of *Klebsiella* phages in general; several have characterised depolymerases with solved structures. The problem is fit. A cocktail assembled from the published collection is matched to a capsule distribution that is not South Asia’s. Phages for South Asian patients will mostly have to be isolated in South Asia, against South Asian strains. Work of this kind has begun: a group in Haripur published a lytic *Klebsiella* phage isolated from local sewage and active against a K17 strain [7]. The same logic is being applied at hospital scale elsewhere — a ventilator-associated pneumonia pilot built its cocktail from retrospective genomic analysis of the unit’s own isolates and reported eradication in 6 of 7 patients receiving the matched cocktail against 0 of 7 receiving a deliberately mismatched one [1]. That contrast is the clinical form of the argument made here.

KL81 is the most defensible single target this analysis identifies. It is South Asian, uncovered, and carbapenemase-positive in every Pakistani genome examined — though that last observation rests on one BioProject and requires replication. A group with access to hospital sewage and a KL81 isolate could close this gap.

The clonality analysis deserves emphasis beyond this dataset. Six of eleven nominally significant geographic differences disappeared when each BioProject was counted once. Comparative work on public bacterial genomes is common, and outbreak-derived genomes are the bulk of what is deposited, so a study that does not test for this will report some clonal expansion as geography. The check costs little.

What this study cannot do is estimate prevalence. Two BioProjects supply most of the Pakistani cohort, and Pakistan has few deposited genomes relative to its burden. Systematic sequencing from independent Pakistani centres would answer the frequency question this dataset cannot. Extending the same analysis across South Asia, where roughly 3000 genomes meet these criteria, would also have the power to resolve the types that collapsed here.

### Limitations

Non-independence rather than sample size is the dominant limitation; only KL81 and KL107 survive Bonferroni correction after collapsing. With effective sizes of 140 and 212, roughly four independent observations against zero are required to reach *P* < 0.05, so types below about 3% frequency in South Asia cannot be resolved; KL36 and KL48 gained no observations when the comparison set grew from 435 to 475 genomes, so more genomes sampled the same way will not resolve them. “No published phage” means none found in PubMed under the searches described, so the uncovered fraction is an upper bound; preprints, non-English literature and unpublished collections were not searched. Both cohorts describe deposited genomes rather than circulating populations. Serotype (K64) and genomic K locus (KL64) were treated as equivalent when reconciling literature to genotypes, which is conventional but not strictly exact. All predictions are computational; no phage was tested against any isolate.

## Conclusion

About half of Pakistani clinical *K. pneumoniae* isolates belong to capsule types with no published phage. At least one, KL81, is South Asian and absent from the regions where *Klebsiella* phages are isolated. Closing that gap requires isolating phages locally rather than importing cocktails matched to another region’s capsule distribution.

## Abbreviations

CRKP: carbapenem-resistant *Klebsiella pneumoniae*
K locus: capsule biosynthesis locus
KpSC: *Klebsiella pneumoniae* species complex
MDR: multidrug-resistant

## Supplementary material

- **Table S1**. Pakistani cohort — 254 accessions with metadata and capsule calls.
- **Table S2**. Comparison cohort — 475 accessions with country group and capsule calls.
- **Table S3**. Curated phage–capsule evidence table — 34 records with evidence grade, PMID and DOI.
- **Table S4**. Full coverage-gap table — all 66 capsule types with burden, carbapenemase count and evidence grade.
- **Table S5**. Independence analysis — raw and BioProject-collapsed counts and *P*-values per capsule type.

## Acknowledgements

The authors thank the research groups whose public genome deposits and published phage characterisations made this analysis possible.

## Author contributions

T.A., conceptualisation, methodology, data curation, formal analysis, software, writing – original draft. A.H.S., data curation, validation, investigation, writing – review and editing.

## Ethical statement

No ethical approval was required. This study analysed only publicly available genome assemblies and associated metadata; no human participants, human material or animals were involved, and no new isolates were collected.

## AI-assistance disclosure

The authors used an AI assistant (Claude, Anthropic) for analysis code development, literature search support, and drafting. All analytical decisions, interpretation and final content are the authors’ own, and the authors accept full responsibility for the accuracy and integrity of the work.

